# Uncovering thousands of new HLA antigens and phosphopeptides with deep learning-based sequence-mask-search *de novo* peptide sequencing framework

**DOI:** 10.1101/667527

**Authors:** Korrawe Karunratanakul, Hsin-Yao Tang, David W. Speicher, Ekapol Chuangsuwanich, Sira Sriswasdi

## Abstract

Typical analyses of mass spectrometry data only identify amino acid sequences that exist in reference databases. This restricts the possibility of discovering new peptides such as those that contain uncharacterized mutations or originate from unexpected processing of RNAs and proteins. *De novo* peptide sequencing approaches address this limitation but often suffer from low accuracy and require extensive validation by experts. Here, we develop SMSNet, a deep learning-based hybrid *de novo* peptide sequencing framework that achieves >95% amino acid accuracy while retaining good identification coverage. Applications of SMSNet on landmark proteomics and peptideomics studies reveal over 10,000 previously uncharacterized HLA antigens and phosphopeptides and in conjunction with database-search methods, expand the coverage of peptide identification by almost 30%. The power to accurately identify new peptides of SMSNet would make it an invaluable tool for any future proteomics and peptidomics studies – especially cancer neoantigen discovery and proteome characterization of non-model organisms.

## Introduction

Typical analyses of mass spectrometry-based proteomics and peptidomics data rely on database-search approaches which provide the best known answers and cannot identify unexpected peptides and proteins. In contrast, *de novo* peptide sequencing attempts to determine the amino acid sequences directly from observed mass spectra but at times suffer from low accuracy. The capability of *de novo* approaches to identify new peptides is crucial for studying non-model organisms with incomplete databases^1^, for identifying uncharacterized mutations or polymorphisms, and for discovering peptides derived from complex processing of RNA and proteins, such as proteasome-mediated splicing^2,3^ and translation of canonically non-coding regions of the genome^4^.

Despite advancements in *de novo* peptide sequencing^5^–^8^, utilization of these methods in routine tandem mass spectrometry (MS/MS) data remains challenging. Recently, DeepNovo^5,9^ has shown that deep learning can be effectively applied to the *de novo* peptide sequencing problem, which resulted in clear improvements over its predecessors. Nonetheless, there is still a huge performance gap in the accuracy and number of identified peptides between *de novo* approaches and standard database-search approaches. Key parts of this limitation lie in the nature of peptide MS/MS spectra which are noisy and sometimes lack crucial information. When interpreting an MS/MS spectrum, evidence for certain amino acid sequence comes in the form of a series of observed ions whose sequential mass differences match to masses of specific amino acids within an error threshold. However, most MS/MS spectra do not contain a complete series of ions that would enable definite deduction of every amino acid position within the original sequence. In these cases, without prior information such as a database of expected amino acid sequences, it would be very challenging or even impossible to arrive at the correct answers.

Here, we introduce SMSNet, a hybrid *de novo* peptide sequencing approach which leverages a multi-step Sequence-Mask-Search strategy to address the issues of missing ions in MS/MS spectra. SMSNet adopts the encoder-decoder deep learning architecture that has been widely used in machine translation^10^, essentially formulating peptide sequencing as a language translation problem. At the initial “sequence” step, SMSNet predict the full amino acid sequence as well as positional confidence scores for each input MS/MS spectrum. Then, during the “mask” step, predicted amino acid positions with low confidence scores are converted into mass tags that indicate the total masses of masked amino acids. Finally, in the “search” step, all predictions are compared against an input amino acid sequence database in order to recover the exact amino acid sequences from mass tags.

Application of SMSNet on large-scaled studies of human leukocyte antigen (HLA) peptidomes and epidermal growth factor (EGF)-treated glioblastoma’s phosphoproteomes reveals over 10,000 previously uncharacterized HLA antigens and over 4,000 previously uncharacterized phosphopeptides. SMSNet’s predictions are in almost perfect agreement with results from database searches and exhibit known characteristics of HLA antigens or could be traced to known phosphoproteins and phosphosites. Furthermore, more than 6,000 of newly identified HLA antigens have not been reported in the Immune Epitope Database^11^ and should contribute to the growing interests in neoantigen discovery for immunotherapy. The power to accurately identify new peptides of SMSNet would make it an invaluable tool for any future proteomics and peptidomics studies.

## Results

### SMSNet model training

We acquired a large collection of >26 million anonymized MS/MS spectra from the Proteomics and Metabolomics Facility at The Wistar Institute for developing SMSNet. This dataset consists of about 1 million unique peptides from diverse species (Supplementary Table S1). Four versions of SMSNet were developed based on distinct training datasets for evaluation on external datasets and comparison with DeepNovo^5^. To compare with DeepNovo, we trained both SMSNet and DeepNovo on a high-resolution MS/MS dataset curated by DeepNovo’s authors (1,422,793 spectra) and on a collection of 1,239,045 highest quality spectra each representing a peptide in our dataset (named WISTARCU-MS-BEST, see Methods). For further evaluations, SMSNet was trained on a set of 25,174,942 spectra of unmodified peptides and peptides containing oxidized methionine (named WISTARCU-MS-M) and on spectra in WISTARCU-MS-M dataset plus additional 1,769,033 spectra of peptides containing phosphorylated serine, threonine, or tyrosine (named WISTARCU-MS-P).

SMSNet employs the encoder-decoder architecture which has been widely used in machine translation^10,12–14^ to predict amino acid sequence from input MS/MS spectra sequentially from the N-terminus to the C-terminus of the peptide (Figure 1. The encoder embeds the input MS/MS spectrum into a fixed-length vector representation through multiple 1D-convolutional and feed forward layers. The decoder, consisting of long short-term memory (LSTM) layers, predicts the likelihood distribution of the next amino acid based on the current state of the model and evidence from the corresponding m/z regions in the input spectrum (candidate ion windows in Figure 1). Once the entire amino acid sequence has been predicted, SMSNet adjusts the confidence score for each position in the prediction through feed forward layers (the rescorer in Figure 1). These steps comprise the “sequence” phase of the Sequence-Mask-Search framework. Then, during “mask” phase, predicted amino acid positions whose confidence scores lie below a user-specified cutoff were replaced by mass tags that reflect their combined masses. Here, we set the confidence score cutoff so that the false discovery rate at amino acid level is less than 5%. Finally, during the “search” phase, SMSNet attempts to recover the exact amino acid sequences from masked positions by searching all predictions against a reference amino acid sequence database.

**Figure 1.**
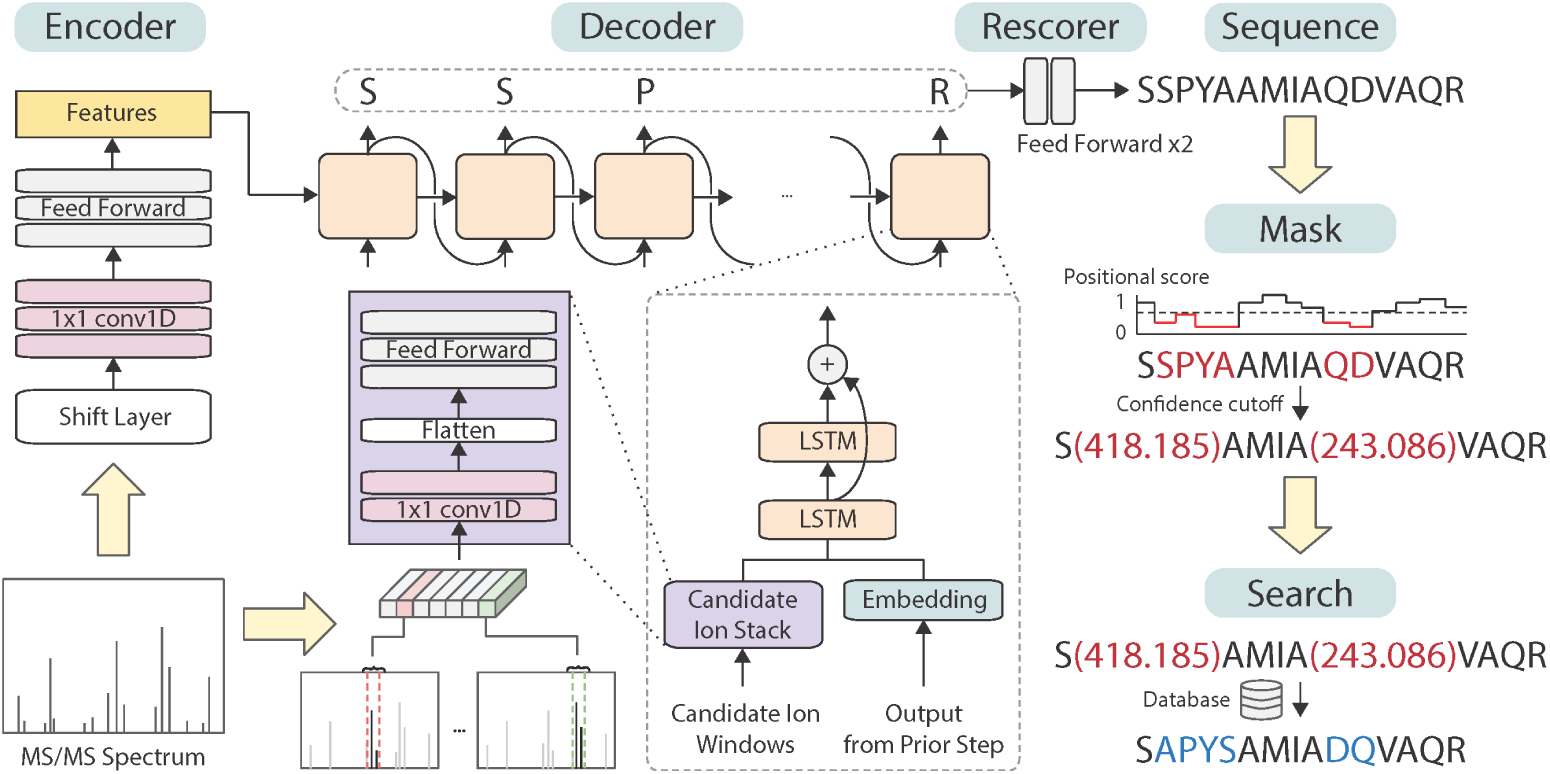
Overview of the Sequence-Mask-Search framework. SMSNet encodes the input MS/MS spectrum and passes the information to the decoder module which outputs amino acid sequentially. During the sequencing process, relevant m/z regions from the input MS/MS spectrum are extracted and fed to the decoder. Post-processing steps involve the adjustment of positional confidence scores, the replacement of low confidence positions by mass tags, and the recovery of exact amino acid sequences in masked segments through database search.

### Performance evaluation on held-out MS/MS data

We evaluated SMSNet’s performance against DeepNovo by training and testing both tools on the same MS/MS spectra from WISTARCU-MS-BEST dataset and from the dataset curated by DeepNovo’s authors. In each test, MS/MS spectra were split so that the train and test sets contain different peptides (Supplementary Table S2). Also, because DeepNovo does not perform any post-processing, outputs from the decoder module of SMSNet prior to score adjustment were used here. At each threshold on the positional confidence score, a predicted amino acid whose score is above the threshold is considered correct only if its mass and prefix mass (the combined mass of earlier amino acid positions in the sequence) differs less than 0.0001 Da and sss0.03 Da from the ground truth, respectively. Predicted amino acids whose scores are lower than the threshold are considered as “unpredicted” and are therefore included in the calculation of recall but excluded from the calculation of precision. At the peptide level, a predicted peptide is considered correct only if all of its amino acid positions whose scores are higher than the threshold are considered correct and that it contains at least 4 predicted amino acid positions. Peptides with less than 4 predicted amino acid positions after applying the score threshold are considered “unpredicted” and treated the same way as “unpredicted” amino acids.

SMSNet consistently outperforms DeepNovo, achieving 21.65% higher amino acid recall than DeepNovo at 5% amino acid false discovery rates on the dataset curated by DeepNovo’s authors (Figure 2a). If all predicted amino acids are considered, SMSNet achieves 71.24% amino acid recall and 47.11% peptide recall while DeepNovo achieves 65.57% and 44.41% recall, respectively. These differences resulted from the fact that SMSNet’s output positional confidence scores are slightly better at distinguishing between correct and incorrect positions than DeepNovo’s do (Figure 2b). Similar difference in performance was observed when both tools were evaluated on our WISTARCU-MS-BEST dataset (Figure 2c). Here, if all predicted amino acids are considered, SMSNet achieves 44.73% amno acid recall and 64.45% peptide recall while DeepNovo achieves 37.57% and 57.02% recall, respectively. Interestingly, the distribution of confidence scores for the correctly predicted positions produced by DeepNovo became bi-modal with a new mode at around 0.6 in this latter test (Figure 2d) while SMSNet’s outputs remain unaffected. We suspect that DeepNovo’s model may have overfit to some coincidental patterns in this dataset. Similar results could be observed when DeepNovo and SMSNet were evaluated on high-quality MS/MS spectra of synthetic peptides^15^ (Figure 2e-f).

**Figure 2.**
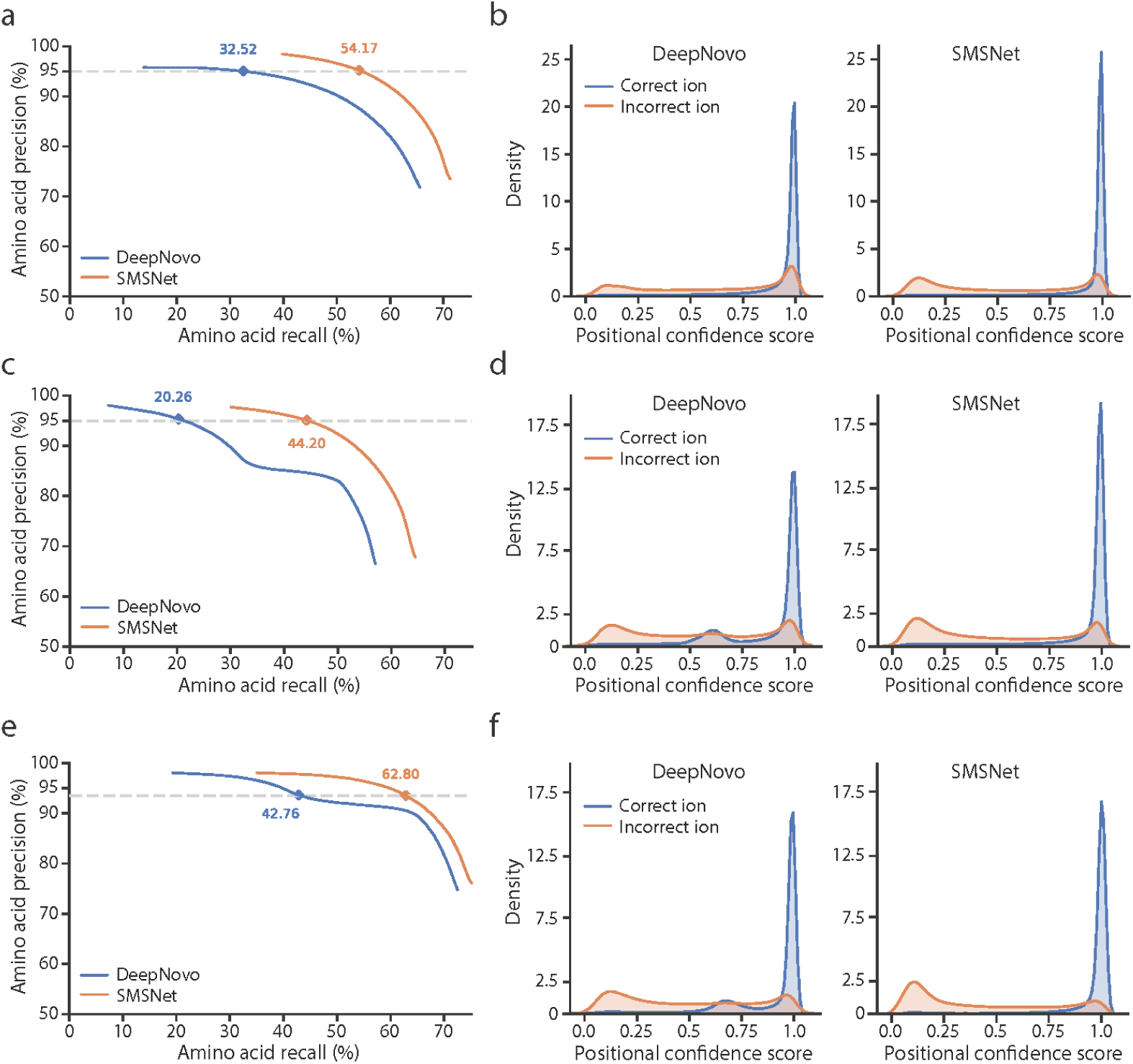
SMSNet outperforms state-of-the-art *de novo* peptide sequencing tool. **a**, Amino acid-level precision-recall curves for SMSNet and DeepNovo when evaluated on the dataset curated by DeepNovo’s authors. The corresponding recalls at 5% amino acid false discovery rate are indicated. **b**, Histograms showing the distributions of positional confidence scores produced by SMSNet and DeepNovo when evaluated on the dataset curated by DeepNovo’s authors^5^. **c-d**, Similar plots showing performances of SMSNet and DeepNovo when evaluated on our WISTARCU-MS-BEST dataset. **e-f**, Similar plots showing performances of SMSNet and DeepNovo when evaluated on high-quality MS/MS spectra of synthetic peptides^15^.

### Impact of the Sequence-Mask-Search framework

By design, the underlying architecture of SMSNet, which sequentially output the likelihood of the next amino acid based on the model’s current state, assumes that the predictions for prior positions are correct and is unable to adjust the confidence scores of prior positions even if later predictions drastically change the context of the sequence. Thus, we trained a neural network module to adjust the resulting positional confidence score based on the information of the whole output amino acid sequence (the rescorer in Figure 1, see Methods). The objective of this rescorer is to maximize the separation in confidence score between correctly predicted and incorrectly predicted amino acid positions. This post-processing step improves the recalls at amino acid level by 9.53% and 9.08% when SMSNet was evaluated on the WISTARCU-MS-M and WISTARCU-MS-P, respectively (Figure 3a-b and Supplementary Figure S1).

**Figure 3.**
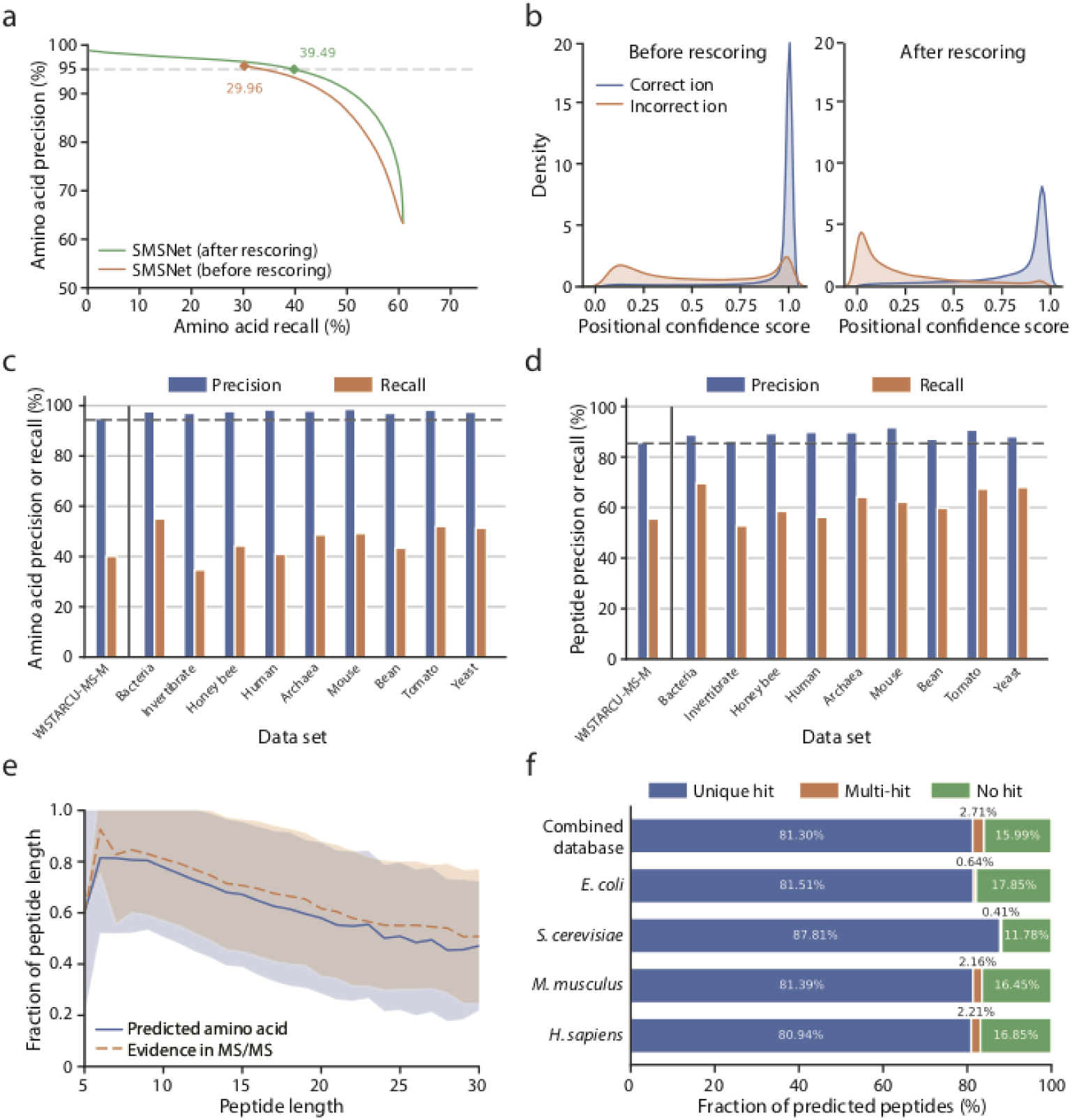
The Sequence-Mask-Search framework significantly improves *de novo* peptide sequencing accuracy. The WISTARCU-MS-M dataset was used to train and evaluate the performance of SMSNet here. **a**, Amino acid-level precision-recall curves for SMSNet before and after the positional confidence score adjustment step. The corresponding recalls at 5% amino acid false discovery rate are indicated. **b**, Histograms showing the distributions of positional confidence scores produced by SMSNet before and after score adjustment. **c**, Bar plots showing amino acid-level precisions and recalls of SMSNet on a test set derived from WISTARCU-MS-M and on MS/MS spectra pf nine species that comprise the dataset curated by DeepNovo’s authors. The threshold on positional confidence score was selected so that 5% amino acid false discovery rate was achieved on the WISTARCU-MS-M test set (the leftmost bars). Dashed line indicate the expected 95% precision based on the applied score threshold. **d**, Similar bar plots showing the results at peptide-level. Dashed line indicate the expected precision level based on the applied score threshold. **e**, Line plots comparing the fraction of predicted amino acid positions that pass the same score threshold used in **c-d** in peptides of various lengths (blue line) to the fraction of amino acid positions that can be definitely determined based on observed ions in the MS/MS spectra (orange dashed line). Shaded area indicate the 1 standard deviation ranges. **f**, Stacked bar plots showing the fraction of predicted peptides that could be matched to various protein sequence databases. Amino acid sequence database for each species was downloaded from Uniprot^16^ (see Methods). Combined database integrates amino acid sequences from all four species considered. In each bar, only predictions whose ground truths exist within the corresponding database were counted. “Unique hit” means that there the predicted sequence matches to exactly one possibility in the database. “Multi-hit” means that the predicted sequence matches to multiple possibilities. “No hit” means the predicted sequence does not match to anything in the database.

Next, we examined the impact of masking predicted positions with low confidence scores on the performance of SMSNet and whether the correct amino acid for each masked position could be recovered by searching against a reference amino acid sequence database. The key concern here is that in order to achieve high accuracy, so many amino acid positions may be masked that the resulting predicted peptides are no longer informative. Here, we trained SMSNet and the rescorer using the WISTARCU-MS-M dataset, identified the adjusted confidence score threshold that corresponds to 5% amino acid false discovery rate on this dataset, and then determined the precision and recall achieved by SMSNet at the same threshold on the dataset curated by DeepNovo’s authors. This revealed that SMSNet with the rescore module performs consistently on MS/MS spectra from diverse species and laboratories at both amino acid and peptide levels (Figure 3c-d). Furthermore, the amount of masked amino acid positions at 5% false discovery rate threshold closely matches the actual number of amino acid positions whose corresponding ions are missing from the corresponding MS/MS spectra (Figure 3e). In other words, SMSNet did not apply too many masks more than the amount required to cover all missing data.

Although the masking step effectively improves the accuracy of SMSNet without too much sacrifice in recall, most utilizations of proteomics and peptidomics data require fully predicted amino acid sequences where mass tags and the Leucine/Isoleucine ambiguity have been resolved. Therefore, we explored whether the correct amino acids that correspond to masked positions could be recovered if an appropriate amino acid sequence database is provided. There are three possible outcomes here. If the predicted peptide is incorrect or contain unexpected amino acid sequence, then there would be no match in the database. On the other hand, if the predicted peptide is correct, but too many masks were introduced, then there may be multiple matches with distinct sequences in the database. Finally, if the predicted peptide is correct and contains a small number of masked positions, a unique sequence hit could be recovered from the database. Our results show that the correct amino acid sequences could be unambiguously recovered for more than 80% of SMSNet’s predictions, even when amino acid sequences from multiple species were used at once (Figure 3f).

### SMSNet discovers more than 10,000 new HLA antigens

To evaluate the utility of SMSNet on real-life mass spectrometry dataset that contains MS/MS spectra of unexpected peptides, we analyzed a large-scale HLA class I peptidome dataset of mono-allelic human B lymphoblastoid cell lines^17^ which consists of more than 35 million MS/MS spectra. The SMSNet model trained on WISTARCU-MS-M dataset was used here because the majority of peptides should not be post-translationally modified. At 5% amino acid level false discovery rate, SMSNet made 95,062 full-sequence predictions and 68,159 partial predictions, which translate to 26,661 peptide-HLA pairs (Fig. 4a). These include 10,702 peptide-HLA pairs not reported in prior study^17^, 8,089 of which are new antigens according to the Immune Epitope Database^11^. Newly identified antigens are of the the right length (8-12 amino acids, Figure 4b) and are also predicted to bind strongly to their corresponding HLA molecules via the correct core sequence motifs (Figure 4c and Supplementary Figure S2). Altogether, these evidences strongly suggest that SMSNet’s predictions are true HLA class I antigens.

**Figure 4.**
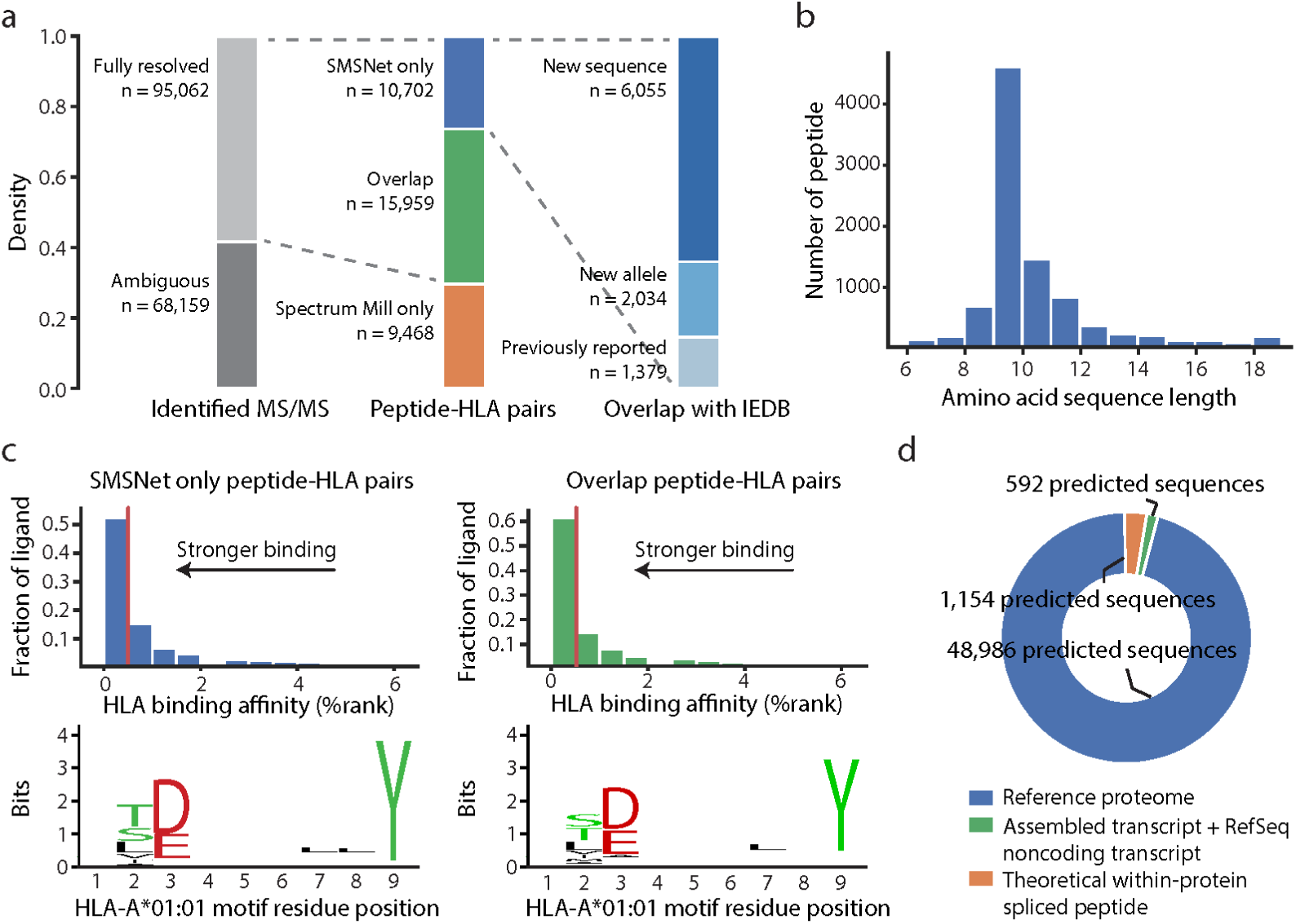
SMSNet uncovers a large number of new HLA antigens. **a**, Stacked bar plots showing the numbers of predictions made by SMSNet and the overlaps between SMSNet and prior study^17^ or the Immune Epitope Database (IEDB)^11^. The number of predicted MS/MS spectra and peptide-HLA pairs are indicated. Comparisons with prior study and IEDB were performed at peptide-HLA pair level. “New sequence” means that the predicted peptide contains amino acid sequence that has not been reported as antigen for any specific HLA allele. “New allele” means that the predicted sequence has been reported to bind to HLA alleles other than the ones considered here. **b**, The length distribution of predicted peptides in 10,702 peptide-HLA pairs newly identified by SMSNet. **c**, Histograms and sequence logos comparing the predicted binding affinities and core sequence motifs between peptide-HLA pairs identified by only SMSNet (left) or by both SMSNet and prior study (right). Binding affinities and core motifs were predicted using NetMHCPan^18^. Vertical red lines designate the 0.5% rank threshold typically used to select strong binders. **d**, Pie chart showing the origins of SMSNet’s predicted peptides. All predictions made by SMSNet were searched against human proteome, transcriptome, and a database of theoretically possible spliced peptides (see Methods).

Additionally, as recent reports indicated that a considerable fraction of HLA antigens may originate from proteasome-mediated peptide splicing^2^, 3 and non-coding regions of our genome4, we explored whether SMSNet discovered any new antigen whose amino acid sequence does not match any human protein in the UniProt database^16^. For this analysis, all predictions made by SMSNet were considered, including those with ambiguous Leucine/Isoleucine and mass tags, in order to determine the origin of as many predictions as possible (see Methods). In total, 48,986 unique predictions directly match to known proteins, 592 unique predictions did not match to known proteins but could be matched to open reading frames in RNA transcripts, and 1,154 unique predictions neither match to known proteins or RNA transcripts but could be explained by splicing of two peptides derived from the same protein (Figure 4e). It should be noted that the number of spliced peptides may be much higher as computational limitation prevents us from considering other possibilities such as inter-protein splicing.

### SMSNet improves the coverage of phosphoproteome

Finally, we evaluated the power of SMSNet to identify post-translationally modified peptides by analyzing a phosphoproteome dataset of control and epidermal growth factor (EGF)-treated glioblastoma cells^19^ that was previously analyzed with MaxQuant^20^. Phosphorylation was selected because of its importance in biology and because our WISTARCU-MS-P dataset contains a sizeable number of phosphorylated peptides (1,769,033 MS/MS spectra) for the model to learn. At 5% amino acid level false discovery rate, SMSNet made full-sequence prediction for 181,144 MS/MS spectra, 134,562 of which are in agreement with previous study (Figure 5a, see Methods). This clearly illustrates the high accuracy of SMSNet that is on par with state-of-the-art database-search approach. Next, we tested whether the addition of SMSNet’s newly predicted sequences to MaxQuant’s search would allow the tool to expand the coverage of its identifications. This results in a net gain of 30,096 identified MS/MS spectra and 3,289 identified phosphopeptides (Figure 5b). In addition to 532 peptides with new amino acid sequences, new identifications also include 2,001 semi-tryptic and non-tryptic peptides which would not be discovered through typical full-tryptic search (Figure 5c). Furthermore, the majority of newly identified phosphopeptides could be observed in multiple replicate samples (Figure 5d) and mapped to known phosphosites and phosphoproteins in the PhosphoSitePlus database^21^(Fig. 5e).

**Figure 5.**
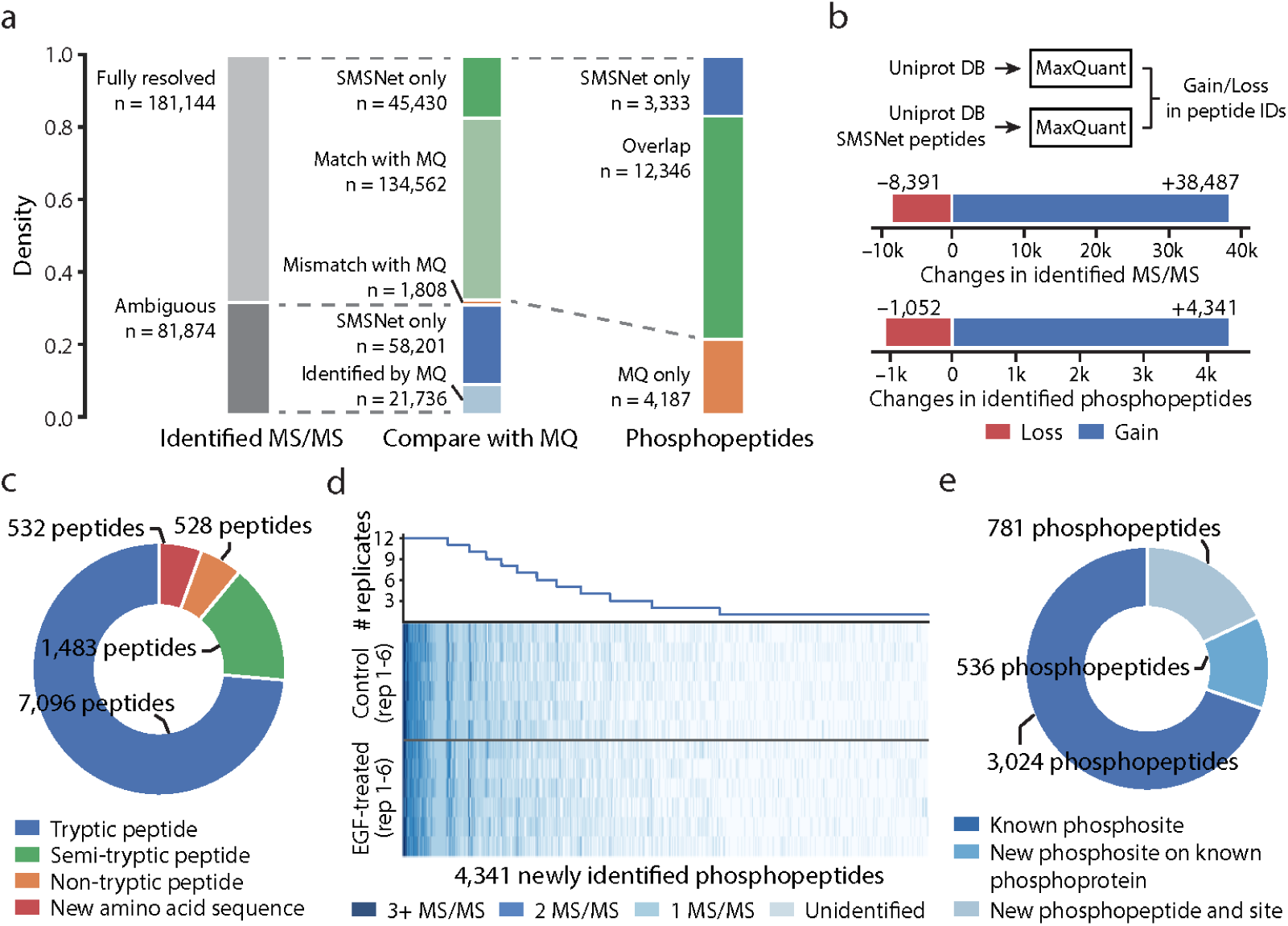
SMSNet improves the coverage of human phosphoproteome. **a**, Stacked bar plots showing the numbers of predictions made by SMSNet and the overlaps between SMSNet and prior study^19^ at MS/MS spectrum and phosphopeptide levels. **b**, The gains and losses in number of identified MS/MS spectra and phosphopeptides after adding SMSNet’s newly identified peptides to the human proteome database and re-analyzing with MaxQuant^20^ (see Methods). **c**, Pie chart showing the composition of newly identified peptides after adding SMSNet’s predictions to MaxQuant’s search. As the search was performed with full-tryptic enzyme specificity, all identifications of semi-tryptic peptides, non-tryptic peptides, and peptides with new amino acid sequences were possible due to sequences supplied by SMSNet. **d**, Heatmap and line plot showing the reproducibility of 4,341 newly identified phosphopeptides after re-analysis using MaxQuant across 6 control and 6 epidermal growth factor (EGF)-treated replicates. Each row in the heatmap corresponds to one mass spectrometry experiment. **e**, Pie chart showing the overlap between newly identified phosphopeptides and known phosphoproteins and phosphosites in the PhosphoSitePlus database2^1^. An identified phosphopeptide was counted as “Known phosphosites” only if all identified phosphorylation sites on that peptide are reported in the database. Identified phosphopeptides that contain unreported phosphosites were grouped based on whether they could be mapped to known phosphoproteins in the database.

## Discussion

SMSNet incorporates several recent computer vision and machine translation techniques^22^–^25^ as well as MS/MS spectra-specific considerations, all of which contribute to its strong performance. The structure of the recurrent neural network in the decoder module was improved from DeepNovo’s design so that more of the relevant information are exposed to the network. To better capture the notion that some amino acid positions are easy to determine from evidences in MS/MS spectra while some could be extremely difficult to do so, we implemented the focal loss^22^. Coincidentally, this feature has also been introduced in a later version of DeepNovo^9^. Interestingly, one of the most impactful features in SMSNet turned out to be the shift layer (Figure 1), which was implemented to help the encoder module identify pairs of MS/MS ions whose mass difference matches to some amino acids. When we removed this layer, the performance of SMSNet dropped by 2.64% and 4.67% at amino acid-level and peptide-level recalls, respectively (Supplementary Table S3). Furthermore, removing the shift filter alone has almost the same effect as removing the whole encoder module. This indicates that incorporating domain-specific knowledge is highly critical for handling complex data generated in the field of biotechnology.

Generalizability is always a major concern in any machine learning study. We have shown that SMSNet not only consistently performs well on MS/MS spectra from diverse species and laboratories (Figure 3c-d) but also is able to identify a large number of HLA antigens that end with non-Lysine/non-Arginine residues even though almost 95% of SMSNet’s training sets are tryptic peptides which end with Lysine or Arginine. Therefore, SMSNet should be able to handle mass spectrometry data generated by any protease enzyme. Additionally, although our evaluations of SMSNet mainly involved MS/MS spectra from Q Exactive mass spectrometers with higher-energy collisional dissociation (HCD), it is possible to adapt the framework of SMSNet for data from different mass spectrometers and peptide fragmentation methods. Part of SMSNet’s capability to adapt to new MS/MS dataset can be seen through its high performance on MS/MS spectra acquired on Orbitrap Fusion Lumos^15^ (Figure 2e).

One limitation of SMSNet is that the effectiveness of the final “search” step, which attempts to recover unique amino acid possibilities for the masked positions, depends on the quality and completeness of the database provided. Although multiple sequence databases could be given to SMSNet at once without apparent deterioration in performance (the combined database in Figure 3f), a large number of predictions remain ambiguous (Figure 4a and 5a). Another possibility is to include all amino acid arrangements that fit the mass tags and perform a scoring-based database search to select the best matches. In this regard, recent advancement in database search using deep learning-assisted prediction of fragment ion intensity^26^ would help improve SMSNet’s “search” step.

Most importantly, we have demonstrated the power of SMSNet to uncover new peptides in large-scale proteomics and peptidomics datasets. High level of agreement between SMSNet and MaxQuant (Figure 5a), together with the findings that SMSNet’s predictions exhibit expected characteristics of HLA antigens (Figure 4b-c) and contain known phosphosites (Figure 5e), indicates that newly identified peptides are true positives. Our results also show that SMSNet can improve the coverage of peptide identification by almost 30% (Figure 4a) and identify new amino acid sequences that better explain observed MS/MS spectra than those in the database do (Supplementary Figure S3). Although many of SMSNet’s new identifications are semi-tryptic and non-tryptic peptides (Figure 5c) which could theoretically be discovered by conventional database-search approach, SMSNet could process 50,000 input MS/MS spectra in 1.14 hours while partial or no enzyme specificity search would take much longer and could result in many false positives. Altogether, SMSNet should become an invaluable tool for future proteomics and peptidomics studies as well as for mining novel peptides in existing datasets.

## Methods

### Data acquisition

A combined mass spectrometry dataset consisting of more than 27 million peptide-spectrum matches (PSM) was obtained from the Proteomics and Metabolomics Core Facility at The Wistar Institute (Philadelphia, PA, USA). All MS/MS spectra were acquired on Q Exactive HF or Q Exactive Plus mass spectrometers (Thermo Fisher Scientific, Bremen, Germany) and processed using MaxQuant^20^ by scientists at the Core Facility. Peptide level false discovery rate was set at 5%. Multiple sets of variable modifications and multiple protein databases were used depending on the goals and scopes of individual mass spectrometry experiments. A partial list of species and the corresponding numbers of peptides identified in this dataset is included in Supplementary Table S1. Importantly, all metadata have been removed to safeguard the identity of principal investigators and the details of their research projects.

From these 27 million PSMs, we constructed three individual training datasets: (i) WISTARCU-MS-M, which consists of 25,174,942 MS/MS spectra that correspond to unmodified peptides and peptides containing oxidized Methionine, (ii) WISTARCU-MS-P, which consists of 26,943,975 MS/MS spectra that correspond to unmodified peptides, peptides containing oxidized Methionine, and peptides containing phosphorylated Serine, Threonine, or Tyrosine, and (iii) WISTARCU-MS-BEST, which consists of 1,239,045 MS/MS spectra that were assigned the highest quality scores by MaxQuant (the “Score” column in evidence output file) for each unique unmodified peptide and charge state. There is no cutoff on the quality score. In other words, the WISTARCU-MS-BEST dataset contains the highest quality MS/MS spectrum for each unmodified peptide at each charge state.

We also acquired three external datasets for evaluating SMSNet’s performance on peptides from diverse species and on MS/MS data from multiple laboratories. For direct comparison with DeepNovo^5^, we downloaded 1,422,793 PSMs from 9 studies of distinct species (PRIDE accessions PXD005025, PXD004948, PXD004325, PXD004565, PXD004536, PXD004947, PXD003868, PXD004467, and PXD004424) that were previously curated by DeepNovo’s authors. For evaluating SMSNet’s ability to discover new peptides, we downloaded 83 raw files consisting of more than 3.5 million MS/MS spectra from an HLA peptidome study of mono-allelic cell lines^17^ (MassIVE accession MSV000080527). Finally, for testing SMSNet-P model’s ability to identify phosphorylated peptides, we downloaded 12 raw files consisting of more than 676,000 MS/MS spectra from a comprehensive phosphoproteome study of control and epidermal growth factor-treated glioblastoma cells^19^ (PRIDE accession PXD009227). High-quality MS/MS spectra of synthetic peptides were acquired from the ProteomeTools HCD Spectral Library^15^. It should be noted that this dataset was acquired on Orbitrap Fusion Lumos mass spectrometer.

### Data preprocessing

MS/MS spectra in the WISTARCU-MS training sets were extracted from raw files and centroided using Thermo Fisher Scientific’s MSFileReader version 3.0. MS/MS spectra in the HLA peptidome and phosphoproteome datasets were extracted from raw files into mgf format using ProteoWizard version 3.0.11133^27^ with the following filter parameters: Peak Picking = Vendor for MS1 and MS2, Zero Samples = Remove for MS2, MS Level = 2-2, and the default Title Maker. Charge state deconvolution was not performed.

For inputting into SMSNet, MS/MS spectra were truncated at 5,000 Da and the observed m/z were discretized at 0.1 Da and Da resolutions to produce vector representations with length of 50,000 and 500,000, respectively. The lower resolution vector provides an overview of the spectrum for the encoder while the higher resolution vector is used by the candidate ion stack. The details of each component are described in the next section.

### Model architecture

Inspired by DeepNovo^5^, we developed our deep learning model focusing on integrating domain knowledge to create a specialized model for *de novo* peptide sequencing, which we called SMSNet. By viewing a peptide sequence as a list of aminoacids, we can view the peptide sequencing problem as a problem of predicting amino acid one by one until termination. Let *X* be an input mass spectrum data, the model can be written as:

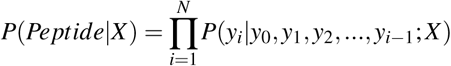

where *y*_*i*_ is the predicted amino acid at position *i, y*_0_ is a special start token, and *N* is the peptide length.

Our model consists of three main components: an encoder, a decoder, and an ion stack. In general, the encoder tries to capture an overview of the input mass spectrum and use it to initialize the decoder. Then, conditioning on the predicted prefix, the ion stack focuses on the relevant part of the spectrum and uses it to compute features for predicting the next amino acid. Finally, the decoder calculates probabilities for the next amino acid using its previous prediction and features from the ion stack. The model architecture is illustrated in Figure 1. Every layers in the networks used rectified linear unit (ReLU) as the activation function unless specified otherwise.

#### Encoder

The encoder was designed to encode an overview of the input spectrum vector into a feature vector of size 1024 which will be used to initialize the hidden state and cell state of the decoder. To integrate the knowledge from the peptide fragmentation process into the model, we restructured the input to make it more likely for the encoder to capture the relationship between positions that could be used to determine amino acid presences. Firstly, the input vector of length 50,000 was duplicated *A* times, where *A* is the number of possible amino acids, into a tensor of shape (50000,A). (*A* is 21 when training on datasets with 20 amino acids plus oxidized Methionines and 24 when training on datasets with 20 amino acids plus oxidized Methionines and phosphorylated Serines, Threonines, and Tyrosines). Each copy of the original input vector is shifted to the left according to each amino acid mass, then padded with zeros. For example, with the resolution of 0.1 Da, the vector representing Alanine is shifted to the left by *floor*(71.037 × 10) = 710. The first 710 values in the vector are discarded, and 710 zeros are padded to the right. This process resulted in a tensor of shape (50000*, A*). Secondly, we created another vector of values from 0 to 49,999 to indicate the index of each positions on the spectrum, then normalized it to have zero mean and unit variance. The index vector was then concatenated to the input to provide the information regarding the position, resulting in a tensor of shape(50000*, A* + 1).

The restructured input was then passed to the encoder neural networks consisting of three 1×1 convolution layers, followed by three fully connected layers. Each of the 1×1 convolution layers applied the same transformation to every input position separately and compute features along the second dimension of the input tensor. This forces the encoder to learn about the structure at each location. The three kernels had shape (1, 32), (1, 64), and (1, 2) that would produce a tensor of shape (50000, 32), (50000, 64), and (50000, 2) respectively after each layer. After that, the feature vector was flatten and passed through three fully connected layers with dimension 512, 512, and 1024, finally resulting in a vector of size 1,024. For regularization, a dropout layer with dropout rate of 0.4 was used between the first and second fully connected layer.

#### Decoder

The decoder is a type of recurrent neural network that receives the feature vector from the encoder and uses it to generated a sequence of amino acids by outputting amino acids one by one. This is similar to the technique used in training neural networks for image captioning^13^ or machine translation^10, 14, 28^ where the input information (an image or a sentence in one language) is encoded into a vector representation, then passed to a decoder to generate the intended output (a caption or a sentence in a different language). Normally, the decoder for image captioning takes only the previously outputted word as input. In SMSNet, the decoder also takes as input a feature vector calculated by the candidate ion stack based on previous predictions for each step. This additional input was designed to provide the model more context about the next amino acid.

In the decoder, we used two layers of long short-term memory (LSTM) of size 512 with layer normalization^24^ on top of each layer and a residual connection^23^ around the second layer. The same encoded vector of length 1,024 was split into two halves and used as initial values for the hidden state and memory in both layers. At each step, the LSTMs take as input a vector of length 544, a concatenated vector between a feature vector of length 512 from the candidate ion stack and an embedding vector of size 32 of the previous amino acid. Then, the output from LSTMs is passed through a fully connected layer with a softmax activation function to produce probabilities for each amino acid. The shape of the last output depends on the number of possible amino acids (20, 21, or 24 depending on the number of modified amino acids considered)

#### Candidate ion stack

Given the total mass of the previously predicted amino acids, the candidate ion stack retrieved relevant sections of the mass spectrum to compute a feature vector for the decoder. Specifically, it looks for evidence supporting the next prediction by focusing on regions that can be the next amino acid. For each possible amino acid, 8 ion types were considered: b, b(2+), b-H2O, b-NH3, y, y(2+), y-H2O, and y-NH3. Suppose there are 21 different amino acids, for each of the 8 ions, we sliced a small window of size 0.2 Da (corresponding to 20 elements at 0.01 resolution) from the original input vector of size 500,000, resulting in 168 20-element vectors. These vectors were stacked together to form an input of shape (168, 20).

The candidate ion stack consisted of two 1×1 convolution layers followed by two fully-connected layers. The idea is to force the model to first learn the peak patterns of each ion, then learn the relationship between ions based on the calculated features. The two 1×1 convolution layers had 32 and 64 filters respectively, while both fully connected layers had 512 dimensions. The output feature tensor was then used as input for the decoder.

### Inference

During inference, we used beam search with beam size of 20 to explore and find the most likely sequence of amino acids. At each step, every remaining hypotheses are ranked by the following formulas^25^:

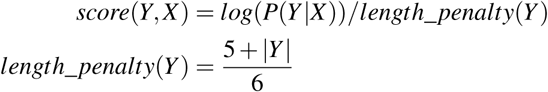

where *P*(*Y|X)* is a product of the previously predicted amino acid probabilities. The length penalty term is used to compensate for longer sequences which usually have lower scores value than the shorter ones. Additionally, during each step, we filtered out hypotheses that the difference between its current mass and the precursor mass did not match any possible amino acid combinations using the knapsack search algorithm.

The beam search decoding would continue until a special ending token is produced or a maximum length of 50 is reached for every remaining beam. After the decoding process ended, the amino acid sequence with the best score according to the provided formulas was selected as the final output.

### Training, validation, and test sets partitioning

To ensure that training, validation, and testing sets do not share a common peptide, we first partitioned unique peptides into three sets, then constructed training, validation, and testing sets from mass spectrum data associated with these peptides. Accounting for the fact that some peptides appear in the datasets much more often than the others, we kept only one random data entry per peptide in validation and testing sets. The validation set was used for choosing the model architecture and determining the number of training steps. For WISTARCU-MS-M and WISTARCU-MS-P, we used validation and test sets of size 50,000. During training, peptides with more than 30 amino acids were ignored.

### Model training

We modeled the peptide sequencing task as a series of amino acid predictions where each prediction is a multi-class classification problem. We chose the focal loss^22^, which is a dynamically scaled cross-entropy loss, as a loss function for our model. For binary classification tasks, the focal loss is defined as:

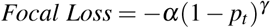

where *p*_*t*_ = *p* for the class with label *y* = 1 and *p*_*t*_ = 1 − *p* otherwise, *p* is the model’s estimated probability for the class with label *y* = 1, *α* and *γ* are hyperparameters for balancing the importance of positive/negative examples and easy/hard examples, respectively. We set *α* to 0.25 and *γ* to 1.0 as it performed best on the validation set. The focal loss is chosen instead of normal cross-entropy loss because we suspected that there is an imbalance between easy examples with complete mass spectrum evidence and hard examples with missing peaks.

To extend the focal loss to multi-class classification, we can view a multi-class classification problem as many binary classification problems. Concretely, we can pass the output of the last layer of the model through multiple sigmoid functions to obtain binary probabilities of being each class, then use the provided formula to calculate the focal loss. For inference, the sigmoid function was substituted with a softmax function to compute probability scores which can be summed to 1.

We initialized all parameters by drawing from a uniform distribution between −0.1 and 0.1, and trained the model using stochastic gradient descent with learning rate decay. An initial learning rate of 0.01 was used until two-thirds of the maximum training step. Afterwards, the learning rate was halved every one-twelfth of the maximum training steps. The gradient of the loss was normalized so that its L2-norm was less than or equal to 5. With a batch size of 32, the models were trained for 4,000,000 steps on WISTARCU-MS-M and WISTARCU-MS-P, which took roughly one month on Nvidia GeForce GTX 1080 Ti.

### Ablation study

To evaluate the impact of each component to the performance of SMSNet, we performed ablation studies by making some modifications to the model, then measuring the performance degradation caused by those modifications. The following modifications were tested:

- Not using layer normalization after LSTM layers in the decoder.
- Using normal cross-entropy loss instead of the focal loss.
- Not considering b-H2O, b-NH3, y-H2O, and y-NH3 ions (neutral loss) in the candidate ion stack.
- Removing the shift mechanism in the encoder. In this variation, we removed the 1×1 convolution layers and fed the low-resolution input vector of size 50,000 directly to the fully-connected layer.
- Removing the encoder entirely and initializing the decoder with a vector of zeros.

Every modified models were trained for 20 epochs on WISTARCU-MS-BEST. The results are shown in Supplementary Table S3.

### Data preprocessing for the rescorer

Unlike the main model, the rescorer model operates solely on the level of amino acids. For each hypothesized amino acid, it predicts the confidence level of the prediction. The following features were used: peptide length, numbers of amino acids with probability more than 0.7, 0.8, and 0.9, a geometric mean of amino acid probabilities in the peptide, the position of the amino acid normalized by the peptide length, probabilities of amino acids at index *t −* 1 to *t* + 2 for current index *t*. We chose these features on the basis that they are not taken into consideration by the main model during the decoding process, and they gave the lowest loss on the validation set. The label for each data point is 1 if the given *de novo* amino acid matches the true label and 0 otherwise.

As the rescorer is designed to evaluate amino acids labels predicted by the main model, we could not used the original training set that the main model was trained on. Therefore, the original validation set was split into rescorer training and validation sets with ratio of a 90:10. The test set were still the same for both tasks.

### Rescorer

We designed the post-processing model to be a shallow neural network consisting of two fully connected layers of size 64. To train the model, we used binary cross-entropy loss and the Adam optimizer with default parameters of *β*_1_ = 0.9 and *β*_2_ = 0.999. The rescoring validation set was used for early stopping.

### Database search for resolving ambiguous predictions

To identify the exact amino acid sequences for predictions that contain mass tags or ambiguous Leucine/Isoleucine positions, we search for possible matches within a given protein sequence database. For analyzing HLA peptidome and human phosphoproteome dataset, the Uniprot^16^ reference human proteome was used. For evaluating whether each of SMSNet’s predictions could be matched to a unique possibility (Figure 3f), the Uniprot reference proteome for either human, mouse (*Mus musculus*), budding yeast (*Saccharomyces cerevisiae*), *Escherichia coli* strain K12, or a combination of all four species was used. All databases include isoforms and predictions proteins. An amino acid sequence within the database is considered a match to an ambiguous prediction if (i) all non-Isoleucine positions in both sequences match, (ii) all Isoleucines in the prediction match to either Leucine or Isoleucine in the database sequence, and (iii) all mass tags in the prediction match to amino acid substrings in the database sequence whose weights differ less than 20 ppm from the corresponding mass tags.

### Comparison with DeepNovo

To compare the performance of our model with DeepNovo^5^, we trained both DeepNovo and SMSNet on two datasets, one from nine species used in DeepNovo^5^ and one from our new dataset, which were used for comparison only. Both models used the same training, validation, and test sets. For DeepNovo, we used the code provided together with their publication.

The first dataset is constructed by combining together all high-resolution datasets in DeepNovo publication. As we only focused on amino acid with Methionine-oxidation, any peptide that contains amino acid with Asparagine- or Glutamine-deamidation in the original dataset was discarded. The remaining data consisted of 1,422,793 mass spectra from 256,200 unique peptides. Due to its lower number of unique peptides, instead of using 50,000 unique peptides as validation and testing sets as in other experiments, we sampled only 20,000 unique peptides from the dataset and used all of their associated spectra, resulting in validation and test sets of size 111,365 and 112,995, respectively.

The second dataset, called WISTARCU-MS-BEST, is a subset of WISTARCU-MS-M dataset that contained only peptide with no amino acid modification. We selected only the spectra with the best quality score according to MaxQuant^20^ for each unique peptide and charge state to form an easy but diverse dataset. In total, there are 1,239,045 spectra of 869,206 unique peptides. The validation and test set each contains 50,000 unique peptide spectra (two spectra were later removed from the test set due to mismatches between their precursor masses and the labels, resulting in the test set of size 49,998). For peptides with many charge states, we randomly chose one charge state and discarded the rest. The summary of all datasets can be found in Supplementary Table S2.

The amino acid vocabulary were set according to the dataset for both models, with 20 possible amino acids for the first dataset and 21 for the second dataset. Apart from the amino acid vocabulary, our model settings were the same as in other experiments. For DeepNovo, we set the spectrum resolution to 0.02 Da and kept other default parameters. At inference time, both model used beam search with beam size 20 to find the most probable peptide for each input.

### Definition of amino acid’s evidence

Given a mass spectrum, we determined that an amino acid has supporting evidence if it follows our defined criteria. Firstly, for an amino acid with mass *M*_*aa*_ and prefix mass *M*_*prefix*_, there must be ions with mass *M*_*prefix*_ and *M*_*prefix*_ + *M*_*aa*_ present in the spectrum. Secondly, a fragmented ion is said to be present in the spectrum if there is at least a peak with any intensity within 0.1

Da of its theoretical b-, b(2+)-, y-, or y(2+)-ion. The first and last amino acids in a peptide only require one ion mass presence.

### Evaluation of HLA-binding affinities and motifs of identified antigens

We downloaded the latest list of known HLA antigens from the Immune Epitope Database (IEDB)^11^ (downloaded April 2019) and searched whether peptide-HLA pairs that were identified by SMSNet have been previously reported. It should be noted that negative entries and entries with ambiguous HLA allele names in the IEDB database were excluded from consideration. NetMHCPan version 4.0^18^ was used to predict the binding affinity (in percentile rank) and the 9-residue core binding motif for each identified peptide-HLA pair. The profiles of allele-specific core binding motifs were visualized using WebLogo^29^.

### Determination of the origins of identified HLA antigens

We iteratively determined the origins of HLA antigens identified by SMSNet by (i) searching all identified amino acid sequences against a reference human proteome downloaded from Uniprot (isoforms and predicted proteins included), (ii) searching the amino acid sequences without any hit in the first step against hypothetical open reading frames extracted from published RNA sequencing data of the cell lines used^17^ and from reference human non-coding RNAs downloaded from RefSeq (GRCh38)^30^, and (iii) searching the amino acid sequences without any hit in the two prior steps against hypothetical spliced peptides generated by joining two peptides from the same protein. The downloaded human proteome includes isoforms and predicted proteins. RNA sequencing data were aligned to the GRCh38 human reference genome using HISAT2. Sequence variants were called using GATK version 4^32^. Reference non-coding RNAs were extracted from GRCh38’s reference transcriptome based on the “ncRNA”, “non-coding RNA”, and “long non-coding RNA” tags. Hypothetical spliced peptides were generated by joining two peptides, each with length at least 3, from non-overlapping regions of the same protein.

### Evaluation of newly identified phosphopeptides

We compared SMSNet’s prediction for each MS/MS spectrum in the dataset to previously reported result^19^ by considering only the actual amino acid sequences without any modification. This is because many predicted peptides contain multiple Serine, Threonine, and Tyrosine residues located near each other and so it is often difficult to pinpoint the exact location of phosphorylation(s). To evaluate the contribution of SMSNet to human phosphoproteome study, we appended newly identified amino acid sequences to a reference human proteome downloaded from Uniprot (isoforms and predicted proteins included) and reanalyzed raw mass spectrometry data using MaxQuant^20^. MaxQuant analysis settings were set as described in the prior study^19^. Namely, variable modifications include Oxidation (M), Acetyl (protein N-term), and Phosphorylation (STY). Enzyme specificity was set as Trypsin/P with 2 maximum missed cleavages. Fixed modification includes only Carbamidomethyl (C). The Match Between Runs and Second Peptide search functionalities were disabled as we want to focus on whether newly identified amino acid sequences would be selected by MaxQuant as the main identification for previously unannotated MS/MS spectra. False discovery rates were set at 1% for both PSM and protein levels. Newly identified phosphopeptides from the re-analysis that were not reported in prior study were searched against the PhosphoSitePlus database^21^ (downloaded April 2019).

### Code availability

The source code and supporting software of SMSNet can be found on GitHub (https://github.com/korrawe/SMSNet). SMSNet was developed under Python 3 and made use of the following freely available packages: Keras version 2.2.4 and Tensorflow version 1.7 (compatible with versions 1.4-1.11).

### Data availability

Processed mass spectrometry data used to train SMSNet (the WISTARCU-MS datasets) will be made available on Figshare upon acceptance of the manuscript. Other datasets used to evaluate SMSNet are already available in public databases and their accession IDs can be found in the Data acquisition subsection in Methods.

## Acknowledgements

This work was supported by the Ratchadapisek Sompoch Endownment Fund, Faculty of Medicine, Chulalongkorn University grant RA62/037 (to S.S.) and the Grant for Special Task Force for Activating Research, Ratchadapisek Sompoch Endowment Fund, Chulalongkorn University (to E.C. and S.S.). We gratefully acknowledge the contribution of mass spectrometry dataset from the Proteomics and Metabolomics Core Facility at The Wistar Institute, the support of the Chulalongkorn Academic Advancement into Its 2nd Century Project, and the donation of TITAN Xp graphic card used in this research by the NVIDIA Corporation. We especially thank Mark A. Knepper (the Epithelial Systems Biology Laboratory, National Heart, Lung, and Blood Institute, National Institute of Health, USA) and Trairak Pisitkul (Systems Biology Center, Chulalongkorn University, Thailand) for facilitating access to computing resources at the National Institute of Health, USA, and for providing critical advice on the manuscript. This work utilized high performance computing resources of the Biowulf cluster, National Institute of Health, USA (http://hpc.nih.gov) and the Center of Excellence for Medical Genomics, Faculty of Medicine, Chulalongkorn University, Thailand.

## Contributions

K.K. developed the software and wrote the manuscript draft. H.-Y.T. and D.W.S. supervised the acquisition and analysis of mass spectrometry data. K.K. and S.S. evaluated software performances. E.C. and S.S. supervised the project. All authors contributed to and approved the final manuscript.

