## Supplementary Tables and Figures for "Uncovering thousands of new HLA antigens and phosphopeptides with deep learning-based sequence-mask-search *de novo* peptide sequencing framework"

### Supplementary Materials

**Supplementary Table S1 – Compositions of species-specific peptides in our WISTARCU-MS dataset**

| Species | Number of Unique Peptides [a] |
|---|---|
| *Homo sapiens* | 165,783 |
| *Mus muculus* | 91,482 |
| *Saccharomyces cerevisiae* | 55,466 |
| *Escherichia coli* | 31,585 |
| *Rattus norvegicus* | 17,587 |
| *Arabidopsis thaliana* | 13,865 |
| *Xenopus laevis* | 11,441 |
| *Bos taurus* | 6,515 |

[a] These numbers include only peptides that could be mapped to exactly one of the databases considered. All databases were downloaded from Uniprot. Isoforms and predicted proteins are included.

**Supplementary Table S2 – MS/MS spectra dataset sizes**

| Dataset | Train | | Validation | | Test | |
|---|---|---|---|---|---|---|
| | Unique Peptide | MS/MS Spectra | Unique Peptide | MS/MS Spectra | Unique Peptide | MS/MS Spectra |
| DeepNovo [a] | 216,200 | 1,198,433 | 20,000 | 111,365 | 20,000 | 112,995 |
| WISTARCU-MS-BEST | 769,208 | 1,095,941 | 50,000 | 50,000 | 49,998 | 49,998 |
| WISTARCU-MS-M | 864,990 | 22,607,416 | 50,000 | 50,000 | 50,000 | 50,000 |
| WISTARCU-MS-P | 994,786 | 24,499,353 | 50,000 | 50,000 | 50,000 | 50,000 |
| ProteomeTools [b] | 162,648 | 212,454 | 20,000 | 26,154 | 20,000 | 26,228 |

[a] MS/MS spectra curated by DeepNovo's authors
[b] High-quality MS/MS spectra of synthetic peptides (ProteomeTools HCD Spectral Library)

**Supplementary Table S3 – Ablation analysis of SMSNet's components**

| Components | Peptide Recall (%) | Difference (%) | Amino Acid Recall (%) | Difference (%) |
|---|---|---|---|---|
| SMSNet (final architecture) | 44.73 | - | 64.45 | - |
| Without layer normalization | 44.10 | -0.63 | 63.89 | -0.56 |
| Cross-entropy loss instead of focal loss | 43.41 | -1.32 | 63.79 | -0.66 |
| Not considering neutral loss in MS/MS | 43.01 | -1.72 | 63.00 | -1.45 |
| Without shift layer in encoder | 40.23 | -4.50 | 61.90 | -2.55 |
| Without entire encoder | 40.06 | -4.67 | 61.81 | -2.64 |

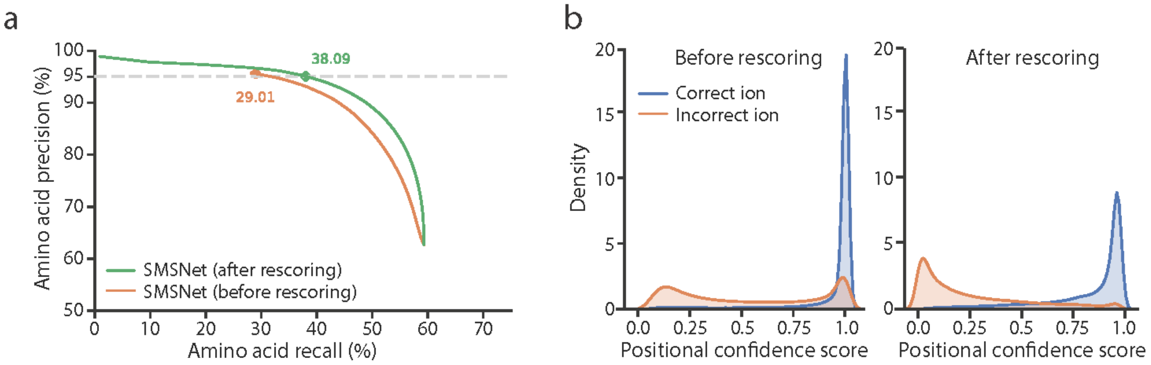


**Supplementary Figure S1 – SMSNet's performance on WISTARCU-MS-P dataset which includes phosphorylation of Serine, Threonine, and Tyrosine. a,** Amino acid-level precision-recall curves for SMSNet before and after the positional confidence score adjustment step. The corresponding recalls at 5\% amino acid false discovery rate are indicated. **b,** Histograms showing the distributions of positional confidence scores produced by SMSNet before and after score adjustment.

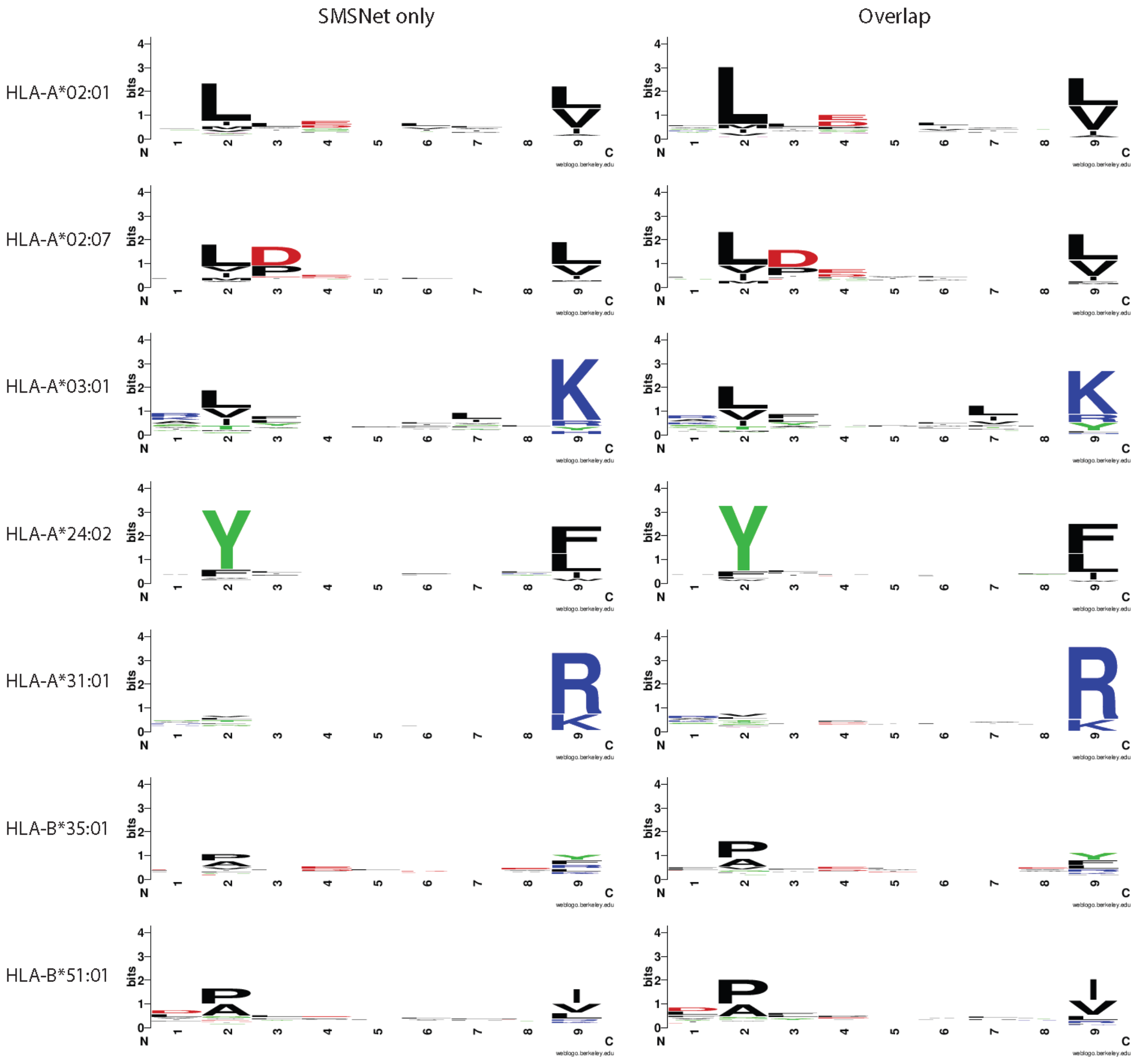


**Supplementary Figure S2 – SMSNet's newly identified HLA ligands possess the same core binding motif as previously reported ligands.** Core binding motifs were predicted using NetMHCPan and the sequence logos were generated using WebLogo. Each row displays the motif for each selected HLA allele. Motifs in column on the left were generated from ligands identified by SMSNet only. Motifs in the column on the right were generated from ligands identified by both SMSNet and the prior study.

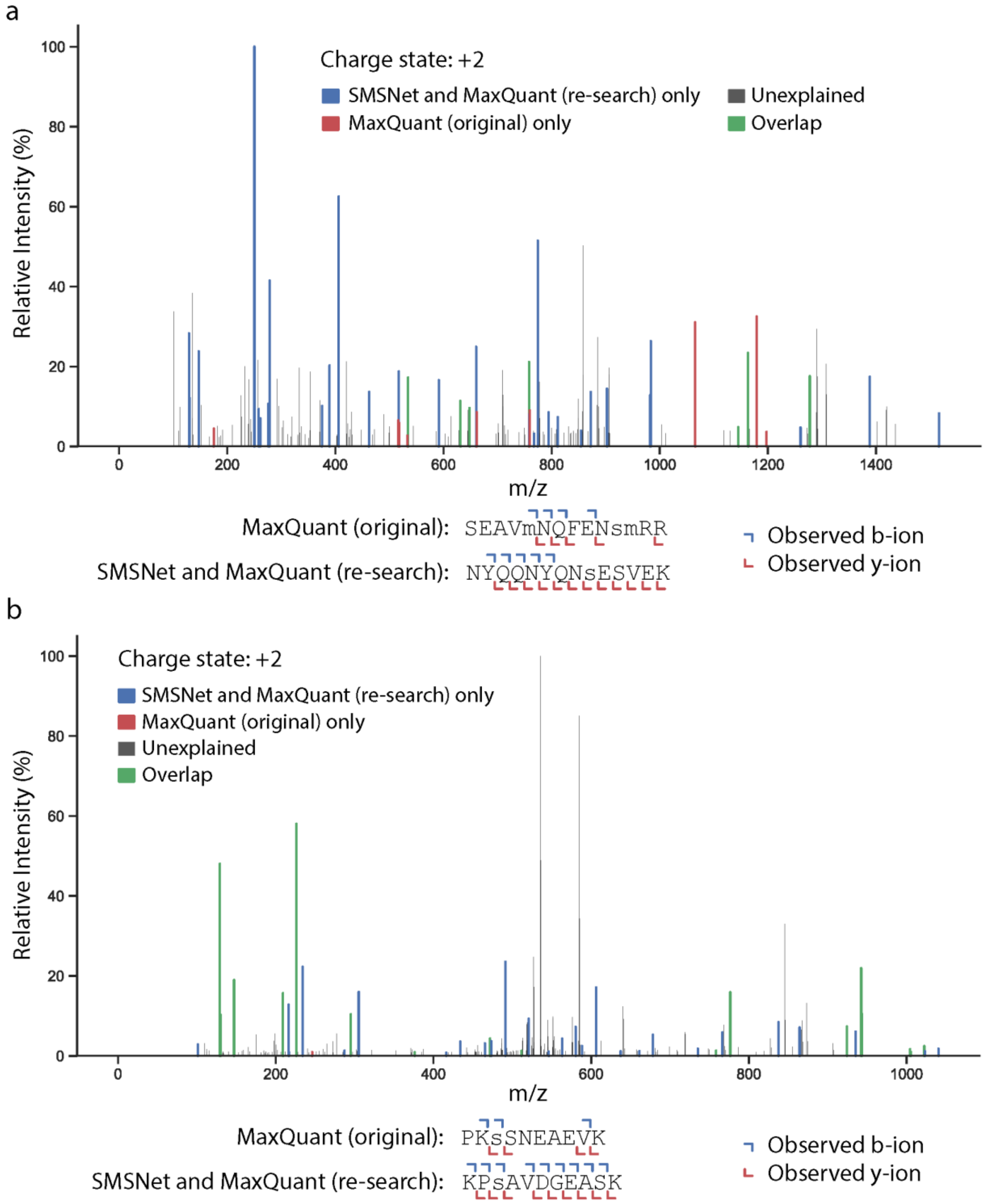


**Supplementary Figure S3 – Examples of annotated MS/MS spectra when SMSNet's and MaxQuant's identified peptides differ.** Amino acids in lower case indicate oxidized Methionine (m) or phosphorylated Serine (s). Observed b-ions and y-ions are indicated in both the sequences and MS/MS spectra.
